# In situ imaging across ecosystems to resolve the fine-scale oceanographic drivers of a globally significant planktonic grazer

**DOI:** 10.1101/2022.08.31.506093

**Authors:** Adam T. Greer, Moritz S. Schmid, Patrick I. Duffy, Kelly L. Robinson, Mark A. Genung, Jessica Y. Luo, Thelma Panaïotis, Christian Briseño-Avena, Marc E. Frischer, Su Sponaugle, Robert K. Cowen

## Abstract

Doliolids are common gelatinous grazers in marine ecosystems around the world and likely influence carbon cycling due to their large population sizes with high growth and excretion rates. Aggregations or blooms of these organisms occur frequently, but they are difficult to measure or predict because doliolids are fragile, under sampled with conventional plankton nets, and can aggregate on fine spatial scales (1-10 m). Moreover, ecological studies typically target a single region or site that does not encompass the range of possible habitats favoring doliolid proliferation. To address these limitations, we combined in situ imaging data from six coastal ecosystems, including the Oregon shelf, northern California, southern California Bight, northern Gulf of Mexico, Straits of Florida, and Mediterranean Sea, to resolve and compare doliolid habitat associations during warm months when environmental gradients are strong and doliolid blooms are frequently documented. Higher ocean temperature was the strongest predictor of elevated doliolid abundances across ecosystems, with additional variance explained by chlorophyll-*a* fluorescence and dissolved oxygen. For marginal seas with a wide range of productivity regimes, the nurse stage tended to comprise a higher proportion of the doliolids when total abundance was low. However, this pattern did not hold in ecosystems with persistent coastal upwelling. The doliolids tended to be most aggregated in oligotrophic systems (Mediterranean and southern California), suggesting that microhabitats within the water column favor proliferation on fine spatial scales. Similar comparative approaches can resolve the realized niche of fast-reproducing marine animals, thus improving predictions for population-level responses to changing oceanographic conditions.

## Introduction

The environmental conditions associated with animal aggregations or population growth can indicate suitable or favorable habitat (Pulliam 2000; Treible et al. 2022). Describing these conditions is a necessary first step towards predicting how populations will respond to environmental change. For most marine animals, however, environmental influences on feeding and reproductive success are not measured at the spatial scales necessary to determine rudimentary physical, chemical, and biological characteristics of their habitat (i.e., the realized niche, Colwell and Rangel 2009). Studies attempting to resolve realized niches are limited spatially (in terms of extent or resolution) or target taxa that do not disperse into all possible habitats (Cadotte and Tucker 2017). To determine the bottom-up components of the realized niche, a target organism should have fast reproduction to overwhelm potential predators (thus limiting top-down impacts) and broad dispersal capability – a life history strategy relatively common in the marine planktonic environment. Given these two biological traits, the small-scale abundances across environmental gradients can enable assessments of the principal bottom-up environmental forces influencing population changes.

Pelagic tunicates are marine animals that epitomize this broad dispersal and fast-reproducing life history strategy. Thus, accurate quantification of their distributions can reveal how the marine environment shapes the realized niche. Salps and doliolids have alternating sexually and asexually reproducing life stages that allow their populations to rapidly increase (Deibel 1998; Walters et al. 2019), sometimes doubling every 8 hours to 1 week (Deibel and Lowen 2012), rivaling maximum population growth rates for any metazoan (Hopcroft and Roff 1995; Paffenhöfer and Gibson 1999). Snapshot distributions of pelagic tunicates, therefore, should indicate recent favorable conditions (Deibel 1998), closely linking the fine-scale aggregations to the realized niche for these taxa (Colwell and Rangel 2009). Aggregations of pelagic tunicates have high clearance rates for phytoplankton and particles (Paffenhöfer et al. 1995; Frischer et al. 2021) and influence biogeochemical cycling through excretion and mass dieoffs (Sweetman et al. 2014; Smith et al. 2016; Luo et al. 2020). Tunicates have the largest predator-to-prey size ratios among marine plankton (Sutherland et al. 2010; Conley et al. 2018) and can display selective feeding (Walters et al. 2019). Because rising average global temperatures are expected to favor smaller plankton (Peter and Sommer 2013), the relative abundance of pelagic tunicates in marine ecosystems may increase (Henschke et al. 2016; Venello et al. 2021; Pinchuk et al. 2021).

Doliolids, in contrast to other pelagic tunicates that can bloom in the open ocean, tend to form dense aggregations in biologically productive shelf- or shelf-break-associated habitats (Lavaniegos and Ohman 2003; Deibel and Lowen 2012). The doliolid life cycle (*Dolioletta gegenbauri*) begins with a fertilized embryo that develops into a larva, oozooid, and then a nurse. This nurse has an anterior extension called a cadophore from which it asexually produces phorozooids that, once released, start asexually producing gonozooids (the only sexually reproducing stage). The phorozooids can release 9-14 gonozooids per day for 8-18 days. At ~20° C, the doliolid life cycle can be completed in 22 days (Paffenhöfer and Köster 2011; Walters et al. 2019). Assuming a single nurse encounters favorable habitat and releases 10 phorozooids, that nurse and the resulting phorozooids can generate 280 gonozooids within 5 days (assuming negligible predation mortality). Thus, fine-scale doliolid abundances can signal conditions favorable for reproduction in the recent past, particularly for *D. gegenbauri*, which can be abundant in subtropical continental shelf ecosystems all over the world (Menard et al. 1997; Paffenhöfer and Gibson 1999; Lavaniegos and Ohman 2003).

Doliolids are frequently detected in continental shelf ecosystems, often associated with upwelling events or offshore intrusions (Deibel 1998; Deibel and Paffenhöfer 2009). Despite their ubiquity, doliolid distributions are not well resolved because the organisms are fragile and difficult to sample quantitatively with conventional plankton nets, particularly on the spatial scales of oceanographic habitat transitions (1-10m, Paffenhöfer et al. 1995; Paffenhöfer and Köster 2011). In situ imaging technologies detect the precise location of individual gelatinous organisms and their associated fine-scale environment, allowing for new descriptions of environmental drivers of aggregations and ecological interactions (Schmid et al. 2020; Robinson et al. 2021; Greer et al. 2021). However, the spatial extent of many of these observations is limited to shipboard measurements that cannot be interpreted beyond the cruise time period (typically a few weeks, Deibel 1998). The realized niche for cosmopolitan animals such as doliolids will remain elusive without observations across spatial scales and ecosystems with different oceanographic regimes.

A comparative approach incorporating observations in contrasting oceanographic environments can advance our understanding of the realized niche for doliolids and similar rapidly reproducing organisms. Resolving these spatial niches has implications for predicting change in marine ecosystem function, as the conditions favoring gelatinous organisms influence the fate of carbon and other biogeochemical processes (Décima et al. 2019; Luo et al. 2020; 2022; Tinta et al. 2021). Existing imagery datasets from different ecosystems during time periods with sharp environmental gradients (summer and early fall seasons) can help determine if common oceanographic variables and vertical water column structure influence doliolid abundance and patchiness in ecosystems with different forcing mechanisms.

## Materials and Methods

### Imaging system and field sampling

All high-resolution images were collected with a towed In Situ Ichthyoplankton Imaging System (ISIIS, Cowen and Guigand 2008). This system has undergone various design iterations, yet the camera setup has had consistent optical properties since approximately 2009. The ISIIS uses shadowgraph lighting and a line-scan camera (Teledyne DALSA) to image water with a 13 cm field of view over a depth of field of 50 cm. The system is towed at a speed of ~2.5 m s^-1^, using motor-controlled wings to undulate between ~1 m from the surface and a maximum depth of ~120 m (or 2-4 m from the benthos) in a “tow-yo” pattern. The camera scans approximately 35,000 pixel lines s^-1^, producing a continuous strip of imaged ocean water that is parsed by acquisition software into 2048 px by 2048 px images (~17 Hz). The ISIIS vehicles used in these studies were equipped with oceanographic sensors, including conductivity, temperature, depth (SBE 49, Seabird Electronics), chlorophyll-*a* fluorescence (ECO FL-RT), and dissolved oxygen (SBE 43) to measure oceanographic conditions associated with each image. The data collected in the Mediterranean Sea had a smaller field of view (10.5 cm), and the vehicle was towed at 2 m s^-1^ with a correspondingly lower scan rate.

Transects were conducted in six different ecosystems and used similar towing methods (Figure 1). The ISIIS tows took place during both day and night, as the image quality is unaffected by ambient light. At least two transects were selected in each ecosystem that were nearest to shore, targeting the known general habitat for doliolids (shelf or near the shelf break). The imagery data were collected off the coast of southern California in October 2010 (Luo et al. 2014), the Mediterranean Sea in July 2013 (Faillettaz et al. 2016), the Straits of Florida in June 2015 (Robinson et al. 2021), the northern Gulf of Mexico in July 2016 (Greer et al. 2021), and northern California and Oregon in July 2018 and 2019 (Briseño-Avena et al. 2020; Swieca et al. 2020).

**Figure 1.**
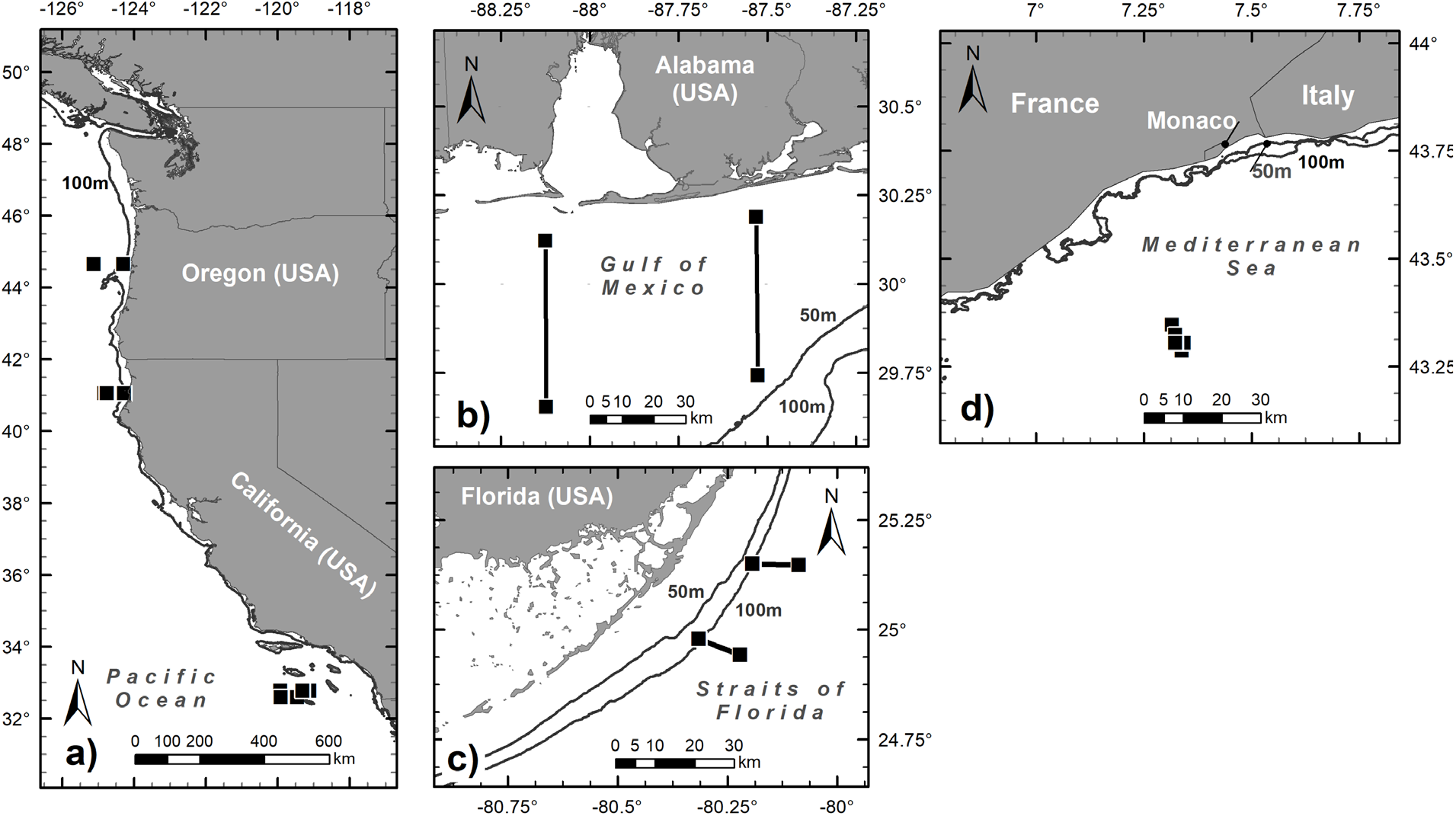
Location of the ISIIS sampling transects conducted within six marine ecosystems. Transects (with total number in parentheses) were analyzed from A) Oregon (2), northern (2) and southern California Current (3), B) the northern Gulf of Mexico (2), C) the Florida Straits (2), and D) the Mediterranean Sea (6).

### Data processing, standardization, and validation

ISIIS data from five ecosystems (all except southern California) were processed using an automated image processing routine that segments the raw images (i.e., extracts regions of interest) and classifies them into >100 taxonomic categories (Luo et al. 2018). A convolutional neural network (CNN) classified the images using different training sets for each ecosystem. This method has produced accurate distributions for many types of zooplankton, including gelatinous organisms and larval fishes (Faillettaz et al. 2016; Schmid et al. 2020; Swieca et al. 2020). All data from southern California was classified manually into one class for doliolids (Luo et al. 2014). While the precise methodological differences in CNN architecture, training, and validation for the different ecosystems are beyond the scope of the work presented here, interested readers should consult the original references that include these details.

Oregon, northern California, northern Gulf of Mexico, and Straits of Florida imagery data from the ISIIS were segmented using k-harmonic means clustering, and classification was performed using a sparse CNN (Luo et al. 2018; Schmid et al. 2021). A probability threshold filtering approach was used to improve precision of the classifier (Failletaz et al. 2016). For Oregon and northern California, precision and recall of doliolid classifications were 82% and 80%, respectively. For the northern Gulf of Mexico automated classification data, model precision was 83% and recall was 93% for all doliolids. The precision and recall was 100% and 7%, respectively, for doliolids in the Straits of Florida. Abundance estimates were derived using a correction factor given these quality control metrics. The doliolids were classified into three life stages (nurse, phorozooid, and gonozooid) in all ecosystems except the Straits of Florida, which classified all doliolids as one category.

For the Mediterranean Sea data, the ISIIS images were segmented using a content-aware object detector (based on a CNN), which ensured an accurate detection of planktonic organisms while ignoring most non-planktonic objects (i.e., marine snow and detritus, Panaïotis et al 2022). Extracted organisms were then automatically classified with a CNN using MobileNetV2 architecture. To improve the precision of the classifier, objects below a 90% probability threshold (established on an independent dataset) were ignored (Faillettaz et al. 2016), resulting in 89% precision and 80% recall for the doliolid class. A subset of these predicted doliolids were imported into the Ecotaxa web application (Picheral et al. 2017) to be sorted into life stage classes (n = 78,193 objects).

To compare data among sampling sites, abundances and associated oceanographic data were summed or averaged over a consistent volume sampled. For organism counts, both stage-specific and total doliolids, were summed across 7-s bins, which corresponded to approximately 1 m^3^ of imaged water. The oceanographic variables were also averaged over this volume, generating a dataset of fine-scale abundances and environmental data for each 1-m^3^ bin. For the Mediterranean dataset, which were processed in a slightly different manner, fine-scale abundances were corrected for discarded images from highly noisy regions around sharp density gradients. Only volumes of water without discarded images were used to calculate patchiness statistics to ensure consistent bin sizes (Bez 2000).

The process of validating data from in situ imaging systems can be arduous and complex when > 60 taxa are considered (Luo et al. 2018; Robinson et al. 2021). For the purposes of this study, we were interested in doliolids and their different life stages: nurses, phorozooids, and gonozooids, all of which are relatively easy to distinguish. Nurses have a long “tail” known as a cadophore, while the gonozooids are a simple barrel-shape. The intermediate stage, both in terms of life cycle and morphological appearance, is the phorozooid, which typically has a disorganized cluster of buds that eventually become free-living gonozooids (Figure 2). For simplicity, we used these three categories to classify the doliolids, but in reality, the gonozooid and phorozooid can be difficult to distinguish if the buds are poorly developed. Likewise, the phrozooid and nurse can resemble one another if the cadophore is poorly developed. The smaller life stages, such as the oozooid, were either discarded or assigned to different doliolid categories (if large enough to be segmented). However, without a training class and associated quality control checks, statistics regarding how these smaller life stages were classified or omitted are unavailable.

**Figure 2.**
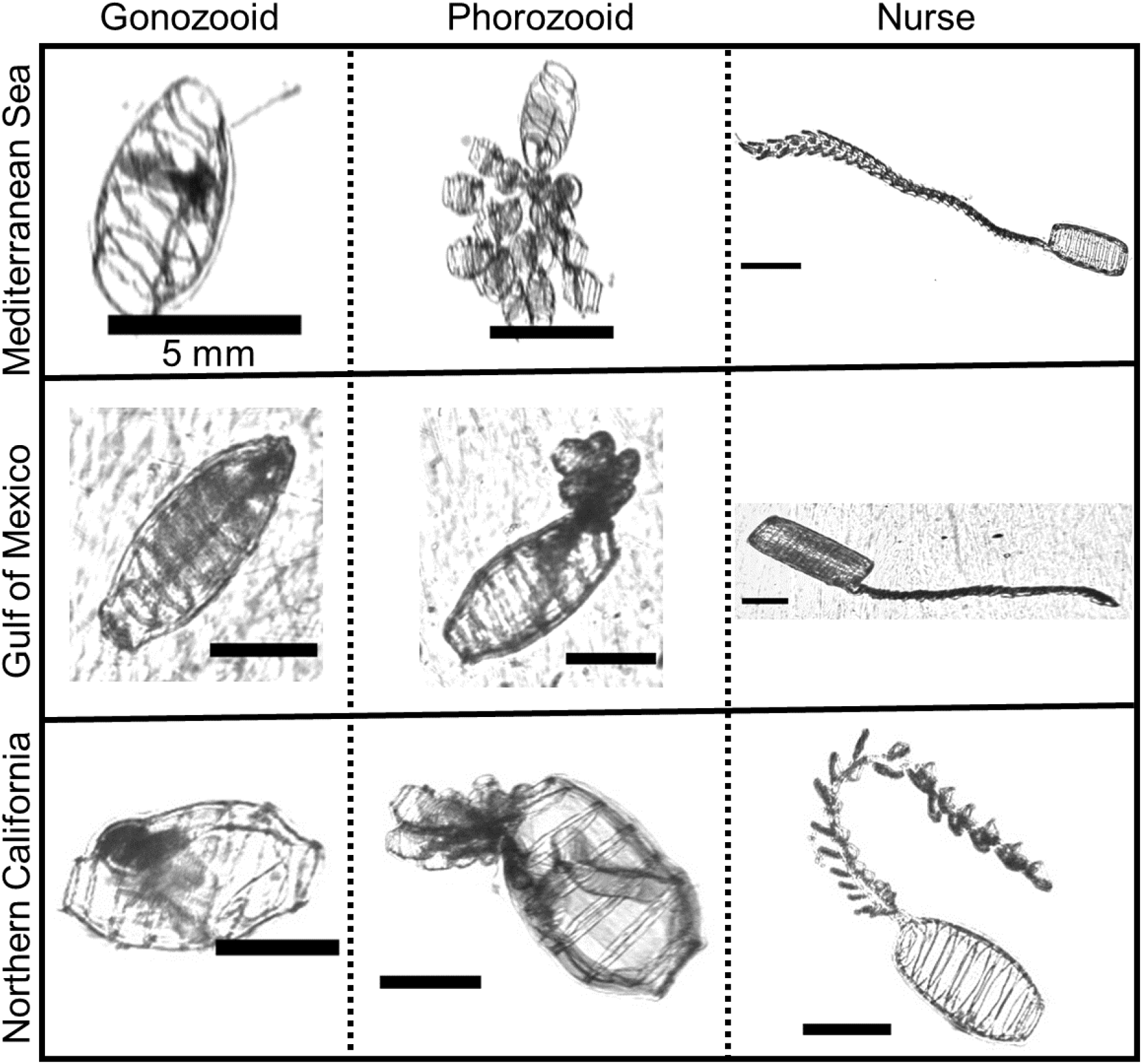
Representative ISIIS images of doliolids from the three different life stages in the Mediterranean Sea, northern Gulf of Mexico, and northern California. All scale bars represent 5 mm.

To evaluate the spatial changes in the accuracy of the automatically generated abundances for these three life stages, we leveraged a human-annotated dataset from the northern Gulf of Mexico. The human-annotated data consisted of regions of interest (ROI) > 3 mm equivalent spherical diameter (Greer et al. 2021) from which “budding” phorozooids, gonozooids (no buds present), and nurses were extracted. This matched the life stages classified in the automated datasets for a more robust comparison. In total, 25,905 doliolids were manually verified from this dataset that encompassed a ~50 km sampling transect just south of Perdido Bay, Florida, USA. These manual identifications were binned using the same procedure as the automated data, and the difference was calculated for each bin and plotted as a histogram to describe discrepancies between automated and manual abundance calculations for each life stage. Spearman rank correlation coefficients were used to quantify the similarity in automated vs. manual distributions among life stages. Discrepancies between manual and automated data could be due to errors in the automated classification or size differences in the water column. Automatically generated abundance data were linearly interpolated for improved distribution visualization for the three life stages (Akima and Gebhardt 2016).

### Mean carbon biomass

We computed the mean carbon biomass of doliolids within the shallowest 100 m for all sites. The average length of doliolids was derived from individual gonozooids in the human-annotated dataset (northern Gulf of Mexico) oriented orthogonally to the camera (aspect ratios > 2.6). The average length of the major axis of the fitted ellipse (8 mm, n = 3,516) was calculated in ImageJ (v1.52a, Schneider et al. 2012) and was used to convert to carbon biomass using the relationship from Gibson and Paffenhöfer (2000):

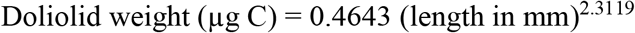

Doliolid carbon biomass m^-3^ was then calculated using the average concentration of individuals in the upper 100 m in each sampling region multiplied by the average individual weight. To place these values in context of previously published Thaliacean carbon biomass data, as well as other mesozooplankton carbon biomass, we compared our values against a data compilation of Thaliacean (doliolids, salps, and pyrosomes combined) and crustacean mesozooplankton biomass (details in Luo et al. 2022). Individual Thaliacean data points were extracted from the Jellyfish Database Initiative (Condon et al. 2015) and averaged to a 1°x1° grid. Crustacean mesozooplankton data were pulled from the NOAA COPEPOD carbon biomass dataset (Moriarty and O’Brien 2013) and averaged to a 1-degree grid. Because an exact location match was not always possible between our sampling sites and the thaliacean and crustacean data compilation, we compared our carbon biomass estimates with the values present in the 2×2 1-degree grids surrounding each site.

We included the crustacean mesozooplankton biomass comparison because gelatinous zooplankton biomass data are very sparse due to poor and inconsistent sampling (Condon et al. 2015). In contrast, crustacean mesozooplankton are sampled with much better spatiotemporal resolution, and a comparison between our estimates of doliolid biomass with crustacean biomass could help to provide better context for their relative abundance and function within ecosystems.

### Statistical analyses and modeling

Data binning, summary statistics, and visualizations were performed in R (R Core Team 2019, v.3.6.1), with extensive use of the packages ggplot2, reshape2, and plyr (Wickham 2016). Patchiness or degree of aggregation was quantified using the Lloyd’s index of patchiness (Bez 2000) on the abundances binned to the same volume with the following equation where x is a set of counts per standard unit volume including zeros:

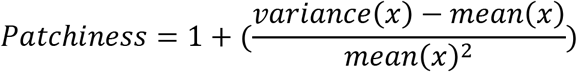

The Lloyd’s patchiness index serves as an indicator of the number of times above the mean abundance each individual organism experiences. For a random distribution, for example, the patchiness index is 1, indicating that an organism will generally encounter the mean abundance. A patchiness index of 100 means that an individual is, on average, experiencing an environment with 100 times the mean abundance (i.e., extremely aggregated). This metric was calculated by re-sampling, without replacement, 20% of the fine-scale abundances in each ecosystem, calculating the patchiness index and repeating 10,000 times to generate a confidence interval. The percent of doliolids that were nurses was calculated for the four environments where life stage-specific data were obtained. These data from each life stage and location combination were fitted with a generalized linear model (GLM) from the quasibinomial family. This GLM family, used to fit proportional data that may contain overdispersion, described the trends in proportions of different life stages compared to fine-scale doliolid abundance. Prior to model fitting, observations above the 99.7 percentile in each ecosystem were removed to reduce the effect of extreme outliers, and from these data, only observations with total doliolids ≥ 15 ind. m^-3^ were included in the model to provide an acceptable sample size for estimated proportions of each life stage.

Structural equation modeling was used to describe the main oceanographic drivers of fine-scale abundances of doliolids across the six ecosystems (n = 21,221). Aggregating across ecosystem-years (‘cross-ecosystem’) and for each specific ecosystem-year, we tested which parameters directly and indirectly predicted doliolid concentrations (individuals m^-3^) using a piecewise structural equation model (PSEM). The predictor variables included depth (m), dissolved oxygen (mg L^-1^), temperature (°C), and salinity. The second-order parameters were fluorescence (relative units, voltage), and water column stratification via the Brunt-Väisälä Frequency (*N*^2^). In the ‘cross-ecosystem’ PSEM, ecosystem-year was included as a discrete random effect (e.g., Oregon 2018) accounting for potential differences in explanatory variable values and doliolid abundances among ecosystems with unknown causes. Data from the Straits of Florida (2014) were omitted from the ‘cross-ecosystem’ PSEM because oxygen measurements were not available.

For all PSEMs, we began with an initial model that reflected the general understanding of causal relationships among variables. However, the initial model included paths with little predictive power and omitted some predictive paths (supp. Figure 1). We added the missing significant paths, and then progressively removed non-significant paths from highest to lowest p-value. Following each removal, we checked a model wide AIC score to confirm that the removal improved model fit. For model visualization and evaluation, we focused on the minimum set of paths that accounted for 90% of the total standardized effect size. PSEMs assign useful p-values and AIC scores to entire causal networks, which include sets of models with a proposed causal structure. Most of the commonly-used models in ecology (e.g., generalized linear models, mixed models) can be incorporated into a PSEM. However, models in the PSEM may not detect highly nonlinear relationships, such as those that could only be captured by a generalized additive model. The goal of this process was to reveal the primary, ecologically relevant relationships rather than extract every statistically significant relationship with the response variable (doliolid abundance).

## Results

### Life stage-specific data validation

Calculated differences between the manually and automatically-derived abundances were life stage dependent for doliolids in the northern Gulf of Mexico (Figure 3a). The gonozooid stage, which had the highest relative abundance in all ecosystems, was frequently detected in higher abundances relative to the manual analysis, as indicated by negative difference values. The nurse and phorozooid life stages, although much less abundant, displayed a normal distribution around zero, indicating the abundances were generally estimated accurately. The distributions from the automated algorithms resembled manually validated life stagespecific patterns for that transect (Greer et al. 2021). The Spearman correlation with the manual analysis was lowest for the gonozooid life stage and was highest for the nurses, which are generally the largest life stage (Figure 3b). These positive correlations, along with quality control checks performed for the different ecosystems, indicated that the automated data could be used to accurately quantify doliolid habitat association patterns across ecosystems. Similar high-resolution snapshot distributions were obtained for the different life stages, or all doliolids pooled, in the other ecosystems.

**Figure 3.**
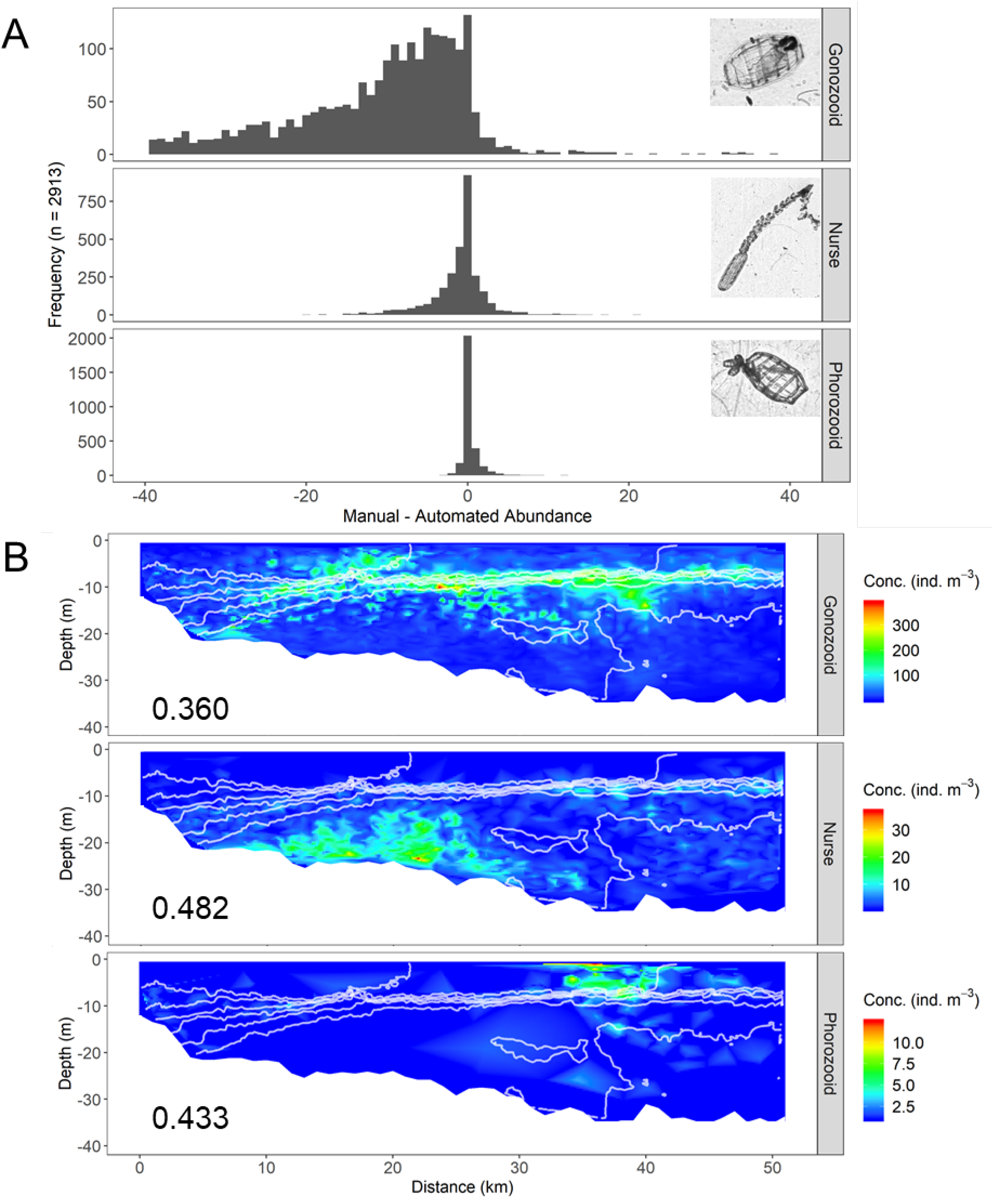
A) Differences between manual and automated fine-scale doliolid abundances in the northern Gulf of Mexico for the three different life stages. B) Example of automated distributions from the eastern transect in the northern Gulf of Mexico for the three different life stages – note changes in the color scale among the three panels. White lines depict salinity contours 30-36. Spearman’s correlation coefficients with the manual data are shown in the bottom left of each panel. Similar fine-scale distributional data were analyzed for all six ecosystems.

### Temperature/salinity and aggregation patterns among ecosystems

Doliolids occupied a wide range of temperatures and salinities and displayed differing abundance patterns among marine ecosystems (Figure 4). The full breadth of the realized niche for doliolids with respect to temperature encompassed 8-32°C, and salinities spanned values from extreme highs found in oligotrophic open oceans (38, Mediterranean Sea) to relatively low salinities typical of river-influenced shelf seas, such as the northern Gulf of Mexico (27). The doliolids were noticeably more abundant at the highest temperatures found in the Mediterranean and the northern California Current (California and Oregon). The northern Gulf of Mexico and Straits of Florida, despite being relatively close both geographically and in temperature/salinity, had remarkably different doliolid abundances, with the former having high abundances throughout the largest temperature and salinity range relative to other ecosystems.

**Figure 4.**
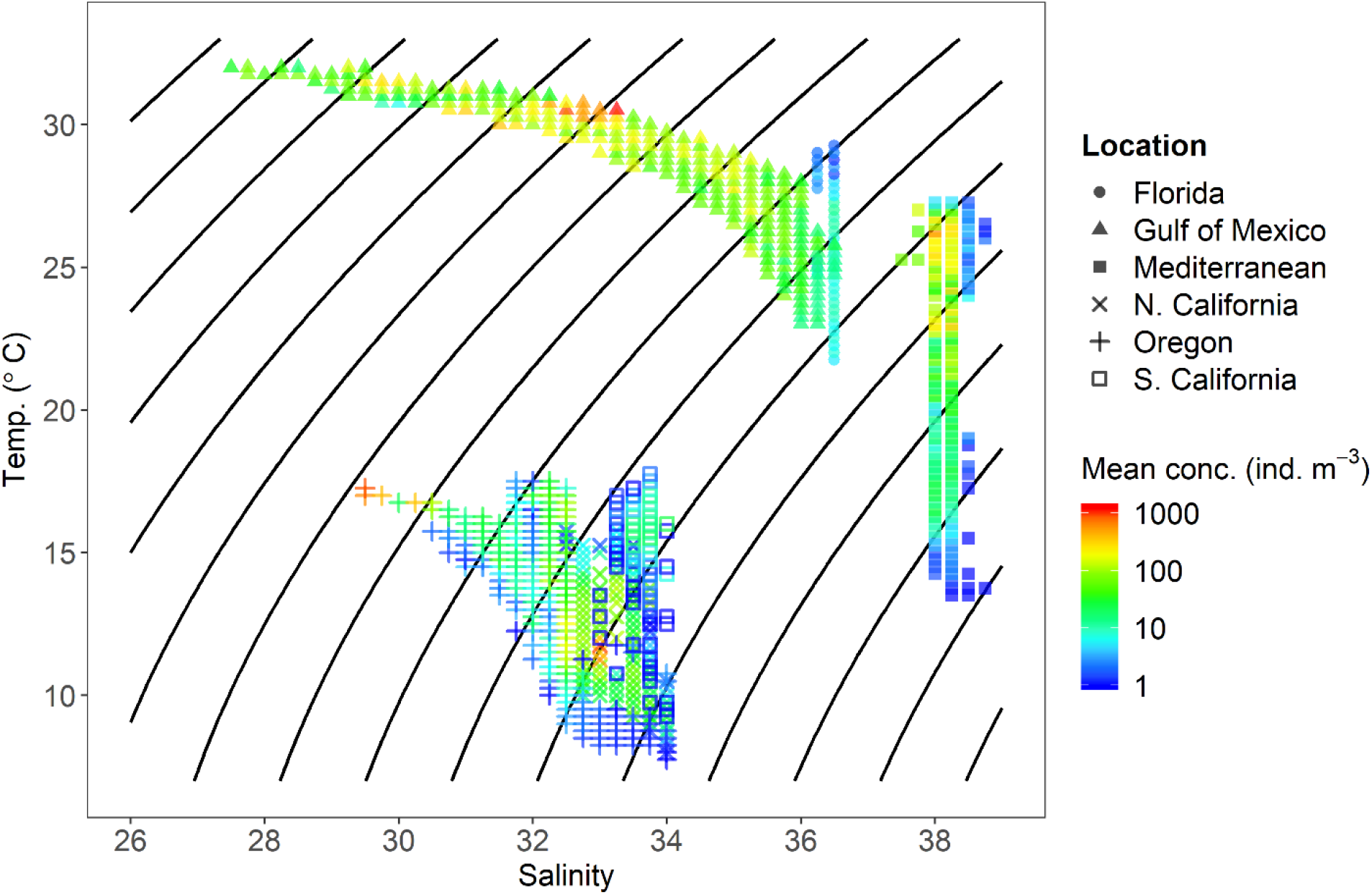
Temperature and salinity for the habitat of all doliolids in different ecosystems. The color of the point corresponds to the mean fine-scale abundance of doliolids quantified in each combination of temperature and salinity (rounded to the nearest 0.25 units). Black lines reference water density contours (σt) 15-30.

Doliolids were generally more aggregated in ecosystems that had lower mean abundances, suggesting they had relatively low abundances in large portions of the water column (Figure 5a). The exception to this trend was the Straits of Florida, which had one of the lowest mean abundances (2.44 ind. m^-3^) and near-random distributions throughout the water column (patchiness of 2.41, random distribution = 1). The other two ecosystems with relatively low patchiness (~10) were the Gulf of Mexico and northern California, which had the highest mean abundances (58.65 and 18.89 ind. m^-3^, respectively). The Mediterranean Sea had the most aggregated doliolid distributions, with Lloyd’s patchiness reaching a mean of ~180, and exhibited more variability relative to the other ecosystems. For the ecosystems with life stage-specific distributions, gonozooids tended to be most abundant and more dispersed compared to the other life stages (Figure 5b).

**Figure 5.**
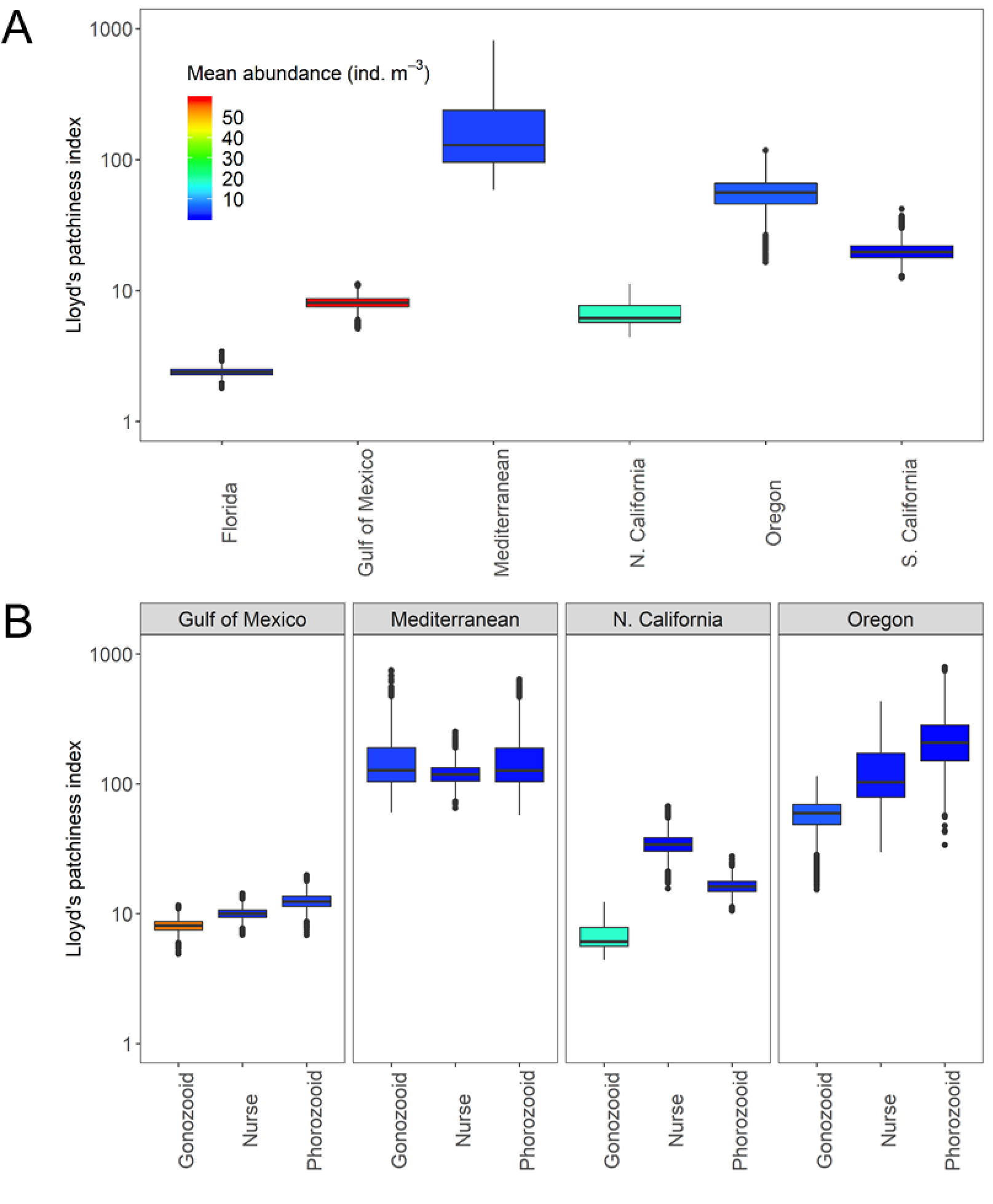
A) Lloyd’s index of patchiness for all doliolids in six different marine ecosystems. B) Life stage-specific patchiness for the four ecosystems where life stages were classified. The abundance color scale is the average across all environments sampled and applies to both panels.

Phorozooids were the least abundant and most aggregated life stage in all ecosystems, with the one exception being northern California where nurses were most aggregated. When the doliolids were not abundant, the relative abundances of nurses increased in the northern Gulf of Mexico and Mediterranean, but the opposite or non-significant trends were found for the two upwelling ecosystems (N. California and Oregon, Figure 6). The Gulf of Mexico was also the only ecosystem where both nurses and phorozooids increased in relative abundance as total doliolid abundance decreased.

**Figure 6.**
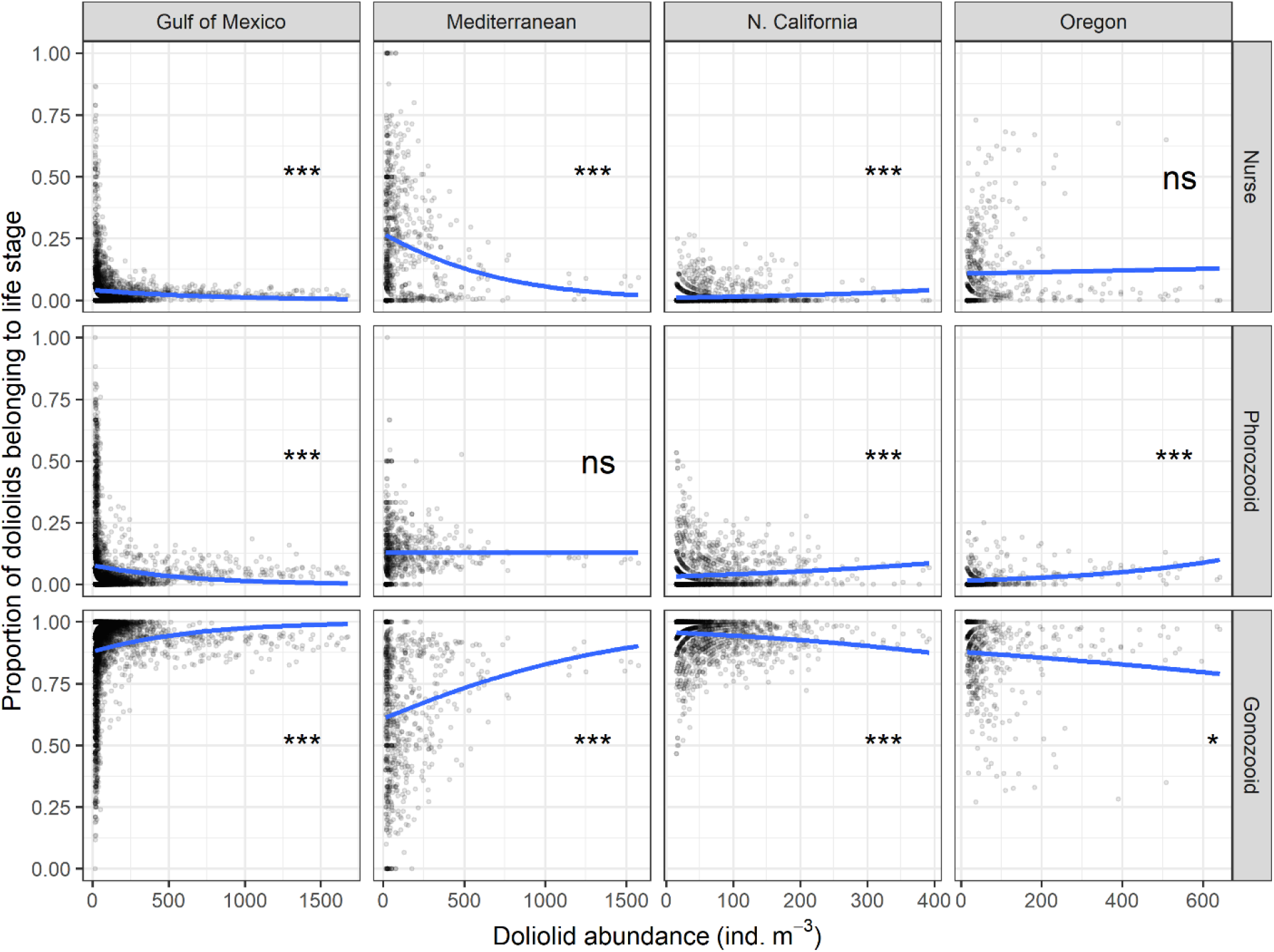
Proportion of the fine-scale abundance belonging to the doliolid nurse, phorozooid, and gonozooid life stages compared to the total doliolid abundance. Four ecosystems are shown where life stage-specific information was available. Blue lines shows the quasibinomial generalized linear model fit to the each subset of data (*** = p < .0001, * = p < .01, ns = not significant). Note that the x-axis scales differ among ecosystems.

### Environmental drivers of doliolid abundances and biomass

Across ecosystems, higher doliolid concentrations were strongly predicted by warmer temperatures (standardized effect size: 0.56). To a lesser extent, doliolids were associated with higher fluorescence (0.22), higher oxygen (0.22), and lower salinities (−0.10). Doliolids were numerically abundant in waters > 20°C with salinities of 28-34 where chlorophyll-*a* fluorescence was elevated. All indirect, predictive pathways for doliolid abundance operated through dissolved oxygen, fluorescence, or both (Figure 7a). Lower fluorescence at depth was linked to greater oxygen concentrations (standard effect size: 0.40). These elevated oxygen concentrations, in turn, were associated with higher doliolid abundances. The degree of water column stratification did not predict doliolid abundance through direct or indirect pathways.

**Figure 7.**
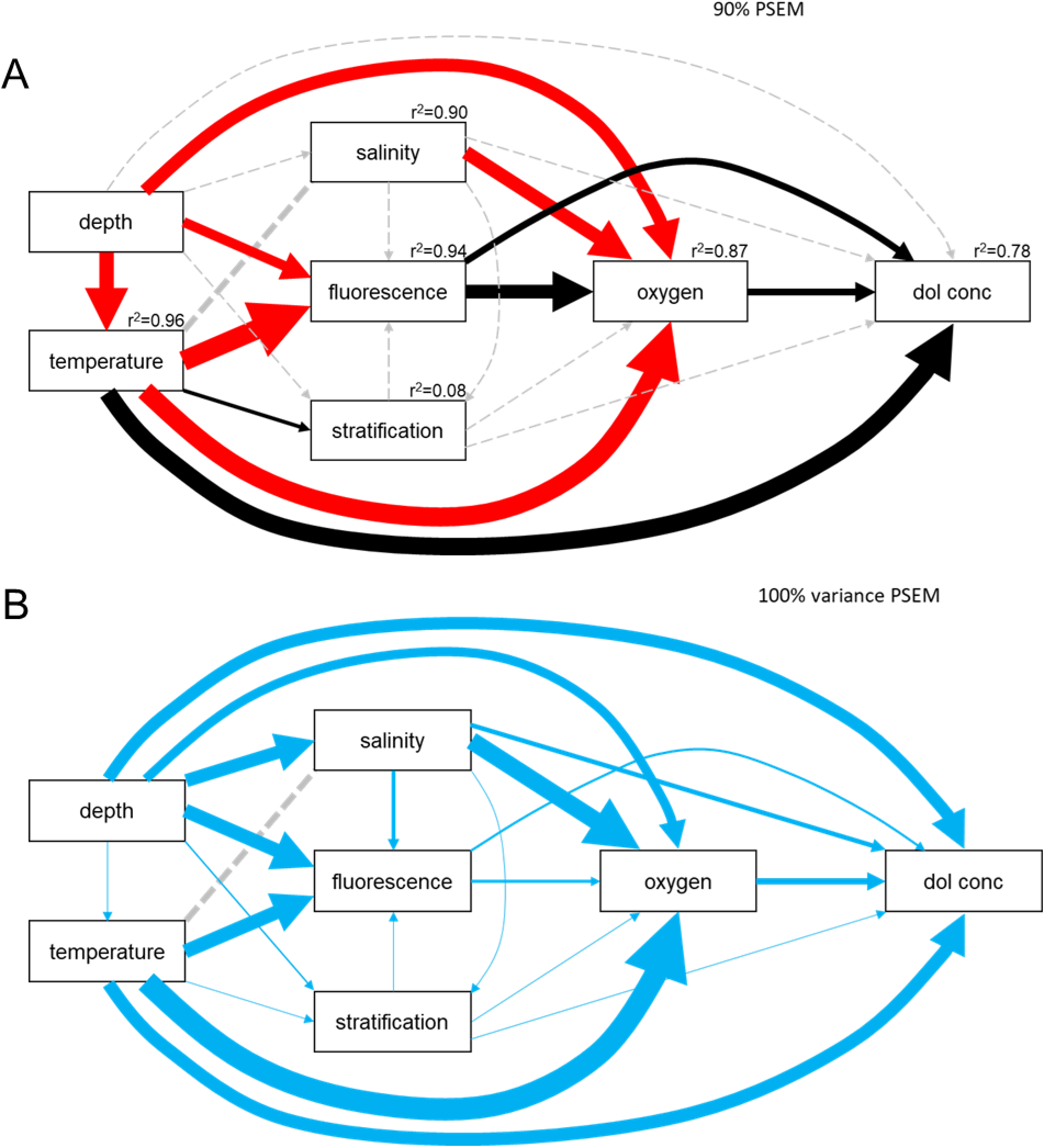
A) ‘Cross-ecosystem’ PSEM with arrows pointing from predictor to response variables. Fluorescence refers to relative chlorophyll-*a* fluorescence. Each arrow’s width is proportional to its standardized effect size. Black and red arrows indicate positive and negative relationships, respectively. The diagram represents the minimum set of paths that account for 90% of total standardized effect size summed across all paths in the model. Dashed gray arrows represent paths included in the initial model but not the minimum set. The total r^2^ for each response variable is displayed above each box. B) PSEM encapsulating 100% of the variance in the doliolid abundance. The arrow width is proportional to the variance of the standardized effect size of each path across ecosystem-years. Narrow arrows indicate the effect was consistent across ecosystem-years, and thicker arrows indicate the effect was variable across ecosystem years. Dashed gray lines represent no causal relationship among the variables in both panels.

Variation in standard effect size of environmental predictors identified which pathways were consistent among ecosystems and years sampled (Figure 7b). Consistent direct predictors of abundant doliolids were salinity, chlorophyll-*a* fluorescence, stratification and dissolved oxygen concentrations. Temperature (variance: 0.27) and depth (variance: 0.25) effects on doliolid numbers were slightly more variable among ecosystem-years. Despite the strong link between temperature and oxygen across ecosystems (Figure 7a), this relationship was highly heterogeneous among ecosystems and years (Figure 7b – see supp. Figure 2 for PSEM differences among ecosystem-years). Conversely, the contribution of temperature to the degree of water column stratification was consistent among ecosystems.

The average length of doliolids from the human-annotated northern Gulf of Mexico dataset was 8 mm, yielding an average individual carbon biomass of 56.8 μg C. The average concentration of individuals from the ISIIS imagery within the top 100 m ranged from 1.35 ind. m^-3^ in Southern California to 58.7 ind. m^-3^ in the northern Gulf of Mexico, yielding mean biomass values from 0.08 – 3.33 mg C m^-3^ (Table 1). These values were compared against the Thaliacean carbon biomass data from the Jelly Database Initiative (JeDI, Condon et al. 2015) and compiled in Luo et al. (2022), as well as the NOAA COPEPOD database (Moriarty and O’Brien 2012). In the grid cells where data were available, doliolid carbon biomass from the present study was lower than thaliacean biomass from JeDI by a factor of 4-10, and lower than crustacean mesozooplankton carbon biomass by a factor of 4-100 (Table 1).

**Table 1.** Mean doliolid biomass (mg C m^-3^) within the top 100 m of the water column at the sampling sites. All sites sampled down to 100 m except for the Gulf of Mexico, where the maximum sampled depth was 35 m. Thaliacean values were compiled from the Condon et al. (2015) JeDI database (see also Luo et al. 2022), and includes salps, doliolids, and pyrosomes. Values are given as the mean biomass in the 2×2 1-degree grid cells surrounding each site where data were present. NA values indicate grid cells with no data.

| Location | Latitude (approx.) | Longitude (approx.) | Mean biomass ($\text{mg C m}^{-3}$) | Thaliacean biomass from JeDI ($\text{mg C m}^{-3}$) | Crustacean biomass from COPEPOD ($\text{mg C m}^{-3}$) |
| --- | --- | --- | --- | --- | --- |
| **Straits of Florida** | 24.9 to 25.2 | -80.3 to -80.1 | 0.14 | 1.45 | 2.53 |
| **N. Gulf of Mexico** | 29.7 to 30.3 | -88.2 to -87.5 | 3.33 | NA | 13.60 |
| **Mediterranean** | 43.3 to 43.4 | 7.2 to 7.3 | 0.21 | 0.84 | 20.00 |
| **N. California** | 41.5 | -124.5 to -124.0 | 1.07 | NA | 16.50 |
| **Oregon** | 44.5 | -125.0 to -124.0 | 0.31 | NA | 8.27 |
| **S. California** | 32.6 to 32.8 | -119.9 to -119.1 | 0.08 | 0.31 | 7.27 |

## Discussion

By using in situ imaging to measure organism abundances during time periods with sharp environmental gradients (warm summer and fall months), we sought to resolve fine-scale habitat characteristics across ecosystems occupied by globally significant planktonic grazers. Doliolids in six marine ecosystems occupied a large range of environmental conditions, and populations were mostly comprised of the gonozooid life stage. The stage-specific fine-scale analysis, however, revealed that nurses became relatively more abundant when doliolids were rare in both eutrophic (Gulf of Mexico) and oligotrophic (Mediterranean) systems, indicating that the nurse life stage may be capable of withstanding unfavorable conditions. The least abundant life stage, the mature phorozooid (with visible buds present), was also more aggregated, indicating that there are micro-habitats favorable for doliolid asexual reproduction in ecosystems with diverse mechanisms of nutrient input, such as riverine input and wind-driven upwelling, or those with scarce nutrients (oligotrophic). The resulting gonozooids can reach high abundances throughout the water column in more eutrophic ecosystems (e.g., northern Gulf of Mexico, northern California). The PSEM indicated that increased doliolid abundances were associated with high temperature and chlorophyll-*a* fluorescence across ecosystems. These findings provide insights into how these abundant, yet understudied gelatinous grazers can thrive circumglobally despite encountering large environmental variations. Furthermore, similar syntheses can be applied to expanding imagery datasets and explanatory variables, generating detailed descriptions of marine organism habitats and ecological niches under changing conditions.

### Scale-dependent patterns and resolving the niche

On large spatial scales, the highest mean abundance of doliolids corresponded to ecosystems with generally higher primary productivity, albeit with different oceanographic forcing mechanisms. The northern Gulf of Mexico receives high amounts of nutrients from various riverine sources (Rabalais and Turner 2019) and was clearly favorable to doliolids even though the images were collected during persistent downwelling conditions (onshore advection of surface waters, Dzwonkowski et al. 2018). In contrast, northern California experiences persistent upwelling-favorable winds during the summer months. At the lower end of the productivity range, the oligotrophic Mediterranean Sea and Straits of Florida had relatively low mean doliolid abundances (2-4 ind. m^-3^). The Mediterranean Sea also supported some of the highest fine-scale abundances in any of the environments, peaking at ~4400 ind. m^-3^, which emphasizes the necessity of measuring on fine spatial scales and resolving corresponding oceanographic characteristics that generate favorable micro-habitats.

Although doliolids are known to associate with the pycnocline and can be abundant within layers containing high diatom concentrations (Paffenhöfer et al. 1987; Greer et al. 2020), cross-ecosystem comparisons revealed that aggregations form in different zones of the water column, not necessarily related to degree of stratification. Doliolids in the Mediterranean Sea were extremely patchy and generally confined to the surface. At the other extreme, conditions in the northern Gulf of Mexico favored high doliolid abundances throughout the water column, with relatively low patchiness and distinct habitat differences among the life stages. Although some studies on the distributions and drivers of doliolid abundances are based on surface expressions only (Ishak et al. 2022), our results demonstrate that the vertical structure of these distributions, even in shallow water columns (< 35 m), is critical to understanding the environments utilized by different life stages and mechanisms for doliolid population sustainability over large spatial scales. In oligotrophic systems, however, the warm near-surface waters may be optimal habitat for doliolids, suggesting that surface samples accurately characterize their peak abundances and environmental drivers.

The suggested positive influence of high temperature and chlorophyll-*a* fluorescence is incomplete without mechanistic linkages through resolving the doliolid prey field. Data from plankton net surveys near Vancouver Island and Alaska (just north of our northernmost study site, Oregon) indicated a strong association of doliolids with recent high temperature anomalies (Venello et al. 2021; Pinchuk et al. 2021), and our results suggest that this can be generalized across ecosystems in terms of fine-scale habitat favorable for doliolids. These high temperature waters had consistently low chlorophyll-*a* fluorescence. Although chlorophyll-*a* fluorescence serves as proxy for phytoplankton abundance, there are caveats with this measurement, such as the source of fluorescence (i.e., phytoplankton), which varies among ecosystems, and the time of day and light intensity where the measurements occur (Carberry et al. 2019). For imaging systems towed over 10s of km, a thorough calibration is often logistically difficult. Non-photochemical quenching can reduce fluorescence, and this effect is enhanced near mid-day, in clearer waters (such as the Mediterranean and Straits of Florida), and at the surface (Sackman et al. 2008; Carberry et al. 2019). Therefore, fluorescence measurements alone are not necessarily representative of the food environment that grazers experience. Laboratory-based experimental studies have shown doliolids are capable of ingesting particles (both fluorescent and non-fluorescent) over a wide size range from <1 μm (bacteria) to >100 μm, with an optimal range of 1–50 μm (Tebeau and Madin 1994; Deibel 1985). However, in nature, larger particles, such as diatom chains over 1 cm in length, can generate high fluorescence and be co-located with doliolid patches (Greer et al. 2020) in mixtures with micro- and nanoplankton. The natural prey field for doliolids may depend on oceanographic processes and plankton size classes difficult to represent in laboratory experiments.

Assessing the full extent of the realized niche for marine animals is often impossible due to issues associated with sampling in terms of accuracy of abundance estimates (poor for fragile organisms), spatial extent (limited to regional studies), and spatial resolution (too coarse to resolve niches in the water column). However, identifying the factors that relate most closely to the realized niche is a critical first step (Chase and Leibold 2003). In more controlled environments, such as enclosed lakes, piecewise structural equation models (PSEMs) have identified top-down and bottom-up controls on zooplankton populations (Du et al. 2015). In contrast, the marine environment is considered a more open system, although physical processes exist that “close” populations and life cycles (Cowen and Sponaugle 2009), and oceanographic variables are often causally linked, influencing biological processes. This presents a challenge because top-down forces (i.e., predation) are dynamic and not well resolved, though evidence suggests these likely influence doliolid abundances (Takahashi et al. 2015) and, more broadly, traits in a variety of zooplankton and phytoplankton (Smetacek 2001; Kiørboe 2008; Grønning and Kiørboe 2020). While it is safe to assume that doliolid reproduction can overwhelm predators within favorable microhabitats, the predation rates relative to reproductive rates of doliolids are largely unknown. More comprehensive sampling and analysis (e.g., other taxa from images and community composition from other sensors), including approaching such data from a PSEM framework, may point to additional explanatory variables and improve our ability to predict community dynamics in different environmental contexts.

### Life stages and bloom dynamics

Although these distributional data represent a snapshot in time, the fast reproductive rates of doliolids due to their multiple asexually reproducing life stages (nurses and phorozooids) provide clues about environmental conditions increasing population fitness on hourly to daily time scales. The sexually reproducing gonozooids are the result of the two asexual phases, so the life stage-specific abundance and composition could indicate a potential bloom (asexual stages) or the aftermath of a recent bloom (gonozooids). Viewing ecosystems on finer scales can reveal dynamic behavior (growth, decay, and all states in between) that is obscured with large scale averaging (Chase and Leibold 2003).

Our data across multiple ecosystems likely encompass a range of growth conditions for doliolids, which may explain the inconsistent relationships between total abundance and proportion of each life stage. Because doliolids are often associated with warm water intrusions (Deibel 1998; Venello et al. 2021), a patchy nurse stage and higher proportion of this stage at high abundances could indicate a recent intrusion. In contrast, the populations in the Mediterranean Sea and Gulf of Mexico may have been more established, as these are marginal seas with potentially less exchange with open oceans compared to relatively narrow upwelling shelves. While this interpretation is somewhat speculative, these results emphasize the necessity of understanding water mass origins and residence times for resolving the niche of planktonic organisms. Furthermore, the cool, nutrient-rich upwelled waters that initiate doliolid blooms (Deibel and Paffenhöfer, 2009), combined with their preference for high temperatures, suggests that there may be a sequence of events, involving upwelling, vertical mixing, and swimming to stay near the surface to generate blooms and patchy distributions. While the timing and spatial details of these events requires further investigation, our results generally agreed with previous studies of life stage-specific distributions, confirming that nurses occur in deeper waters relative to the phorozooid and gonozooid stages, which tend occupy near surface waters and dominate total biomass (Paffenhöfer et al. 1995; Takahashi et al. 2015; Greer et al. 2021).

Because the nurses have an unknown longevity and high reproductive potential (Walters et al. 2019), the survival of this stage is likely critical for doliolids colonizing new habitats in subtropical and temperate shelf ecosystems all over the world. Specifically, the Gulf of Mexico and the South Atlantic Bight are both favorable for doliolid blooms during the summer (Paffenhöfer et al. 1987; Greer et al. 2020; Frischer et al. 2021). There is likely some degree of population connectivity between these two areas via the Loop Current and Gulf Stream (Paffenhöfer and Lee 1987), suggesting that some proportion of doliolids can survive the journey through the potentially less favorable south Florida ecosystem (as indicated by low measured abundances). Evidence across ecosystems suggests that nurses are critical for population dispersal because they tend to encompass a higher proportion of the life stages when total doliolid abundances are low, similar to patterns described in the Kuroshio Current (Takahashi et al. 2015). Nurses are sensitive to hydromechanical signals via sensory cells located on the cadophore, which may aid in avoiding predators that target the relatively nutritious part of their body (Paffenhöfer and Köster 2011). These advanced sensory capabilities could have an added benefit of enabling the nurse to locate portions of the water column that reduce or promote dispersal, depending on oceanographic context. Further study on the swimming behaviors of doliolids, and whether they are life stage or food concentration dependent (Deibel 1998; Frischer et al. 2021), would improve predictions of transport and population dispersal.

### From images to ecological function

In situ images have been collected in different marine environments over the past three decades, with the most impactful findings often related to some key aspect of ecosystem functioning only revealed by accurate and spatially-detailed sampling. For taxa such as the colony-forming cyanobacteria *Trichodesmium*, resolving fine-scale patchiness across ocean basins improved estimates of surface ocean nitrogen fixation (Davis and McGillicuddy 2006). For processes such as carbon and silica cycling, image data have revealed that large, abundant, and historically under-sampled protists (*Rhizaria*) play key roles (Biard et al. 2018; Stukel et al. 2018). The common thread among these studies is that in situ imaging data have direct implications for global-scale marine biogeochemical processes that would have not been resolved without both sampling accuracy and spatial detail.

Connections to biogeochemical processes are not as direct for doliolids and other metazoans. Carbon biomass relative to other zooplankton can serve as proxy for their contribution to biogeochemical fluxes. Doliolid biomass, as estimated from ISIIS imagery sampled in the shallowest 100 m, ranged over an order of magnitude across the various sampling sites even within the same current system (e.g., sites in the California Current). The shallowest environment sampled (northern Gulf of Mexico, 35 m), had the highest biomass concentration at 3.33 mg C m^-3^. While difficult to directly compare against the total Thaliacean biomass data compilation of Luo et al. (2022), which includes salps and pyrosomes averaged across all seasons and in a coarse grid (~100 km horizontal resolution), doliolid carbon biomass values from the ISIIS data were typically ~10-30% of these values. When compared against large crustacean biomass values (Moriarty and O’Brien 2013), the ISIIS-derived doliolid biomass values were approximately 1-2 orders of magnitude lower, as the large crustacean mesozooplankton ranged from 2.53 mg C m^-3^ off the coast of Florida to 20.0 mg C m^-3^ in the northern Mediterranean. The areas with the largest difference in doliolid and crustacean mesozooplankton biomass were the oligotrophic systems (Mediterranean and S. California), whereas the eutrophic northern Gulf of Mexico saw the smallest difference between doliolids and crustacean biomass.

Considering their mean abundances throughout the water column, on a global scale, doliolids may not substantially contribute to carbon and other biogeochemical fluxes compared to crustaceans and more broadly dispersed salps and pyrosomes. However, considering the fine-scale abundances and aggregations of doliolids in a variety of environments, they likely have a local impact on biogeochemical processes, especially in biologically productive shelf environments (e.g., northern Gulf of Mexico), that are yet to be resolved. Considering doliolids’ grazing preferences and high predator-to-prey size ratios (Conley et al. 2018; Walters et al. 2019), they may have a trophic role more similar to the microzooplankton. In contrast to the microzooplankton, which consume between 62-67% of primary production in the global oceans, and up to 77% regionally (Calbet and Landry 2004; Schmoker et al. 2013), the average doliolid biomass and grazing impact is significantly lower. Results from this study suggest that conditions favorable to doliolid aggregations frequently vary, particularly within a water column. These aggregations are capable of consuming 100% of the phytoplankton biomass (Deibel 1998). As opposed to salps, which have fecal pellets that can sink >1500 m d^-1^ (Phillips et al. 2009), doliolid fecal pellets sink relatively slowly (13-107 m d^-1^; Patonai et al. 2011), so their biogeochemical impact may trace water masses where they aggregate and influence the microbial loop. Improved doliolid physiological measurements, abundances, and rate measurements of adjacent biogeochemical processes on fine spatial scales through time (hours – days) will be necessary to assess doliolid contributions to marine food webs.

## Conclusions

We analyzed six high-resolution datasets to resolve bottom up drivers of abundance and aggregation tendencies for a fragile and fast-reproducing marine animal in contrasting oceanographic regimes. This is a first step towards resolving the realized niche for doliolids, as there is rich potential for targeted analyses using a comparative approach and applying machine learning tools to describe, in detail, organism habitats and interactions with predators, competitors, and prey (Schmid et al. 2020; Greer et al. 2021). Similar approaches for other target organisms could reveal the n-dimensional ecological context necessary for the organism to thrive - something that cannot be determined from regional studies alone, or without appropriate sampling technology resolving spatial scales relevant to individuals and their habitat changes (1-10 m). Fine-scale spatial distributions are lacking for most zooplankton, especially within different oceanographic contexts. Resolving processes on these scales is critical because their behavioral and ecological responses may be driven by what they perceive in their immediate environment. Because many gelatinous organisms are understudied yet influential in biogeochemical cycling (Henschke et al. 2016; Luo et al. 2022), further studies within regions and over longer time scales would be valuable for quantifying the ecological influence of gelatinous organisms from bottom-up and top-down perspectives. With improving image analysis techniques, an increasing diversity of sampling platforms (e.g., autonomous vehicles, moored instruments, etc.), and computational infrastructure for data synthesis, similar approaches could be taken to generate a global, high-resolution map of life (Jetz et al. 2012) for marine organisms.

## Acknowledgements

Captains and crews of many different research vessels enabled the successful collection of field data analyzed in this study. These vessels included the R/V *Point Sur*, R/V *Bell M. Shimada*, R/V *Walton Smith*, R/V *Sikuliaq*, R/V *Sally Ride*, R/V *Téthys II*, and the R/V *Atlantis*. Thanks also to Cedric Guigand and Charles Cousin (Bellamare, LLC) for technical support on numerous field expeditions. We thank two anonymous reviewers for their helpful comments, as well as Marco Corrales for providing the NOAA internal review. Funding support was provided by NSF OCE 2023133, NSF OCE 1419987, NSF OCE 1737399, NSF OCE 2125407, NSF XSEDE OCE170012, and the Gulf of Mexico Research Initiative. K.L. Robinson was supported by a grant from the Copyu Foundation. We thank Christopher Sullivan and Dominic Daprano at Oregon State University’s Center for Quantitative and Life Science for computing support. The authors declare no conflicts of interest.

## Supplemental Figures

**Supp. Figure 1.**
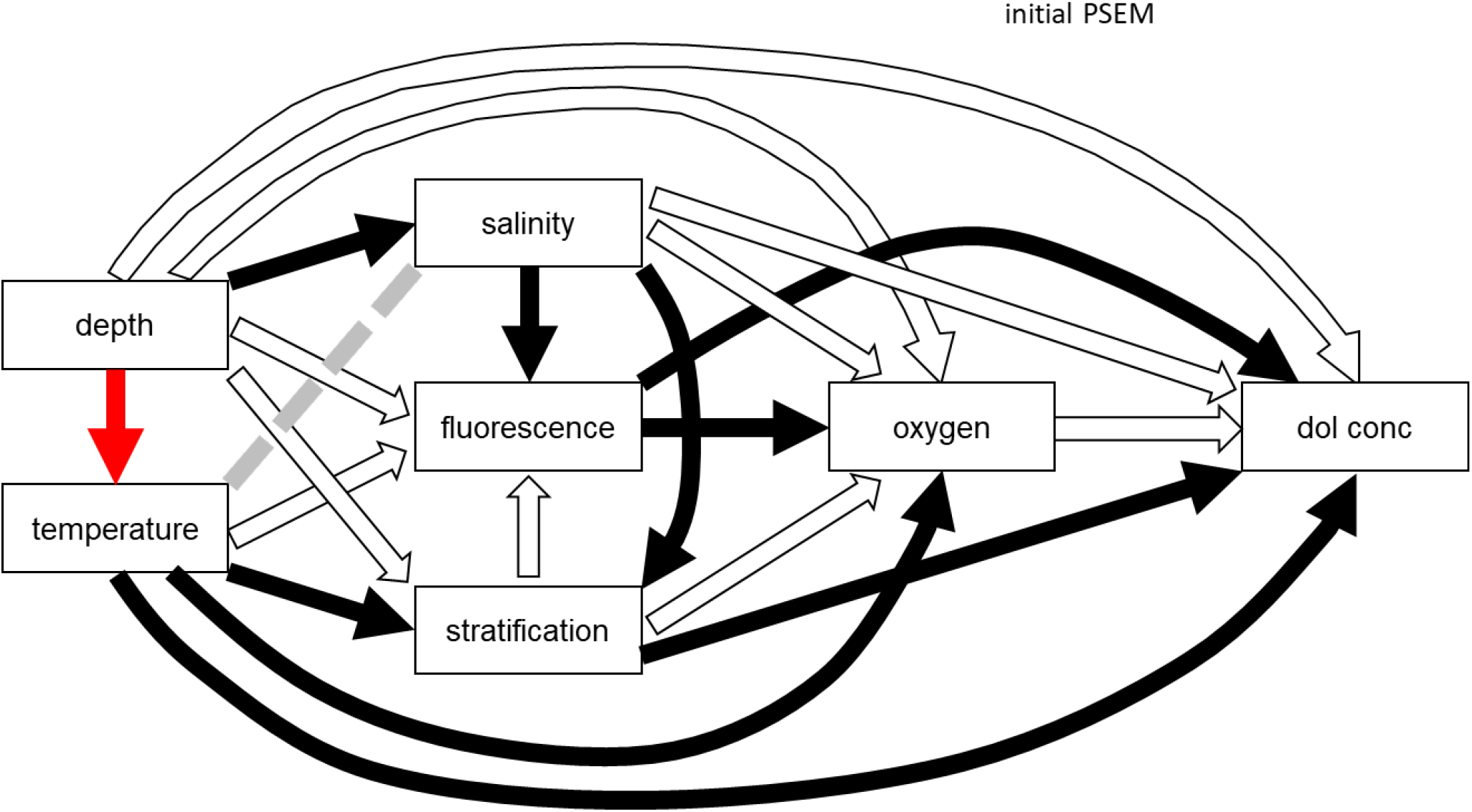
Initial piecewise structural equation model derived from plausible relationships among oceanographic variables. The red lines indicate negative relationships, while the black and white lines indicate positive and neutral relationships, respectively.

**Supp. Figure 2.**
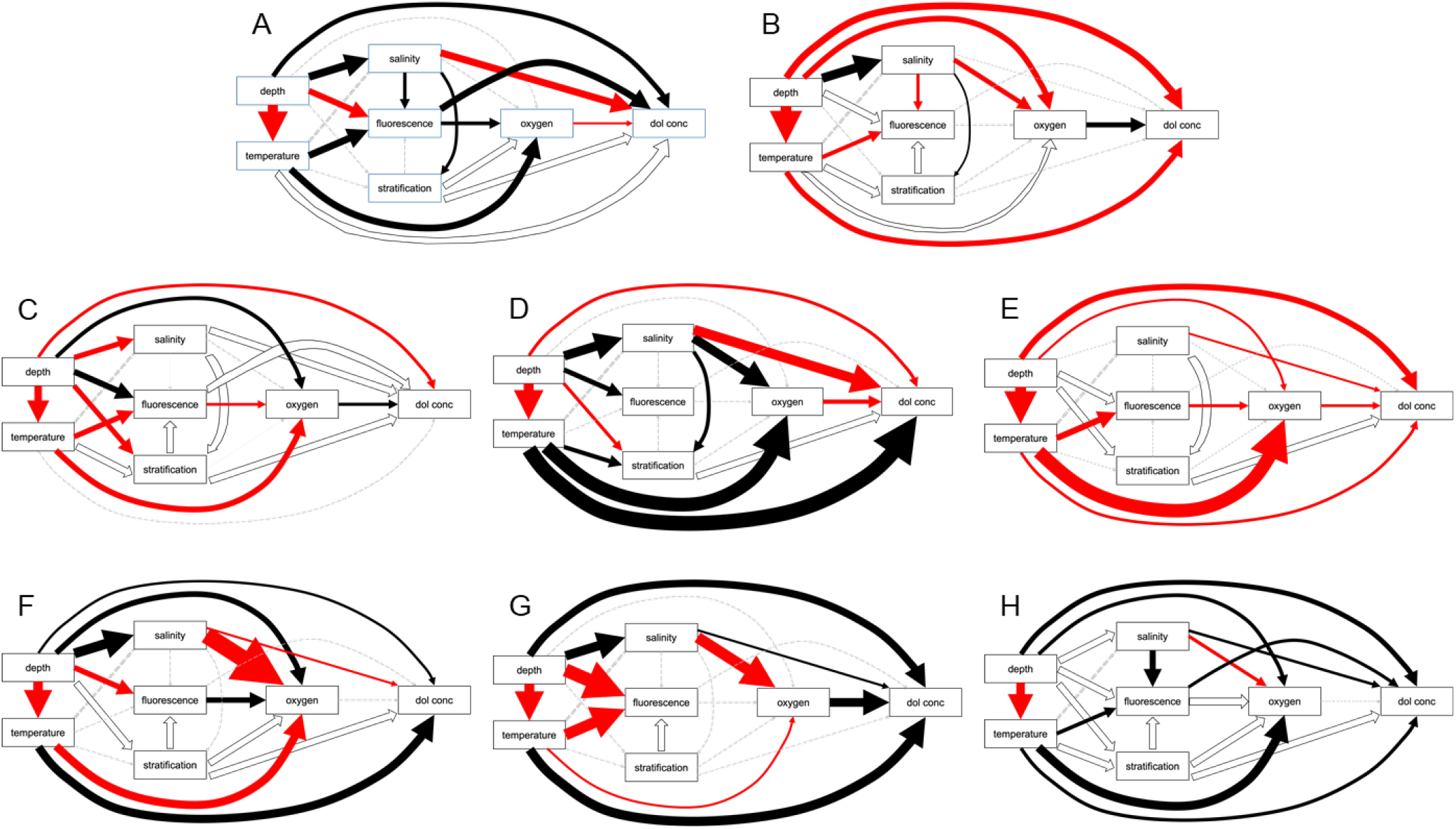
The results of the 90% piecewise structural equation models for the different ecosystem-years in A) N. California 2018, B) N. California 2019, C) Straits of Florida 2015, D) northern Gulf of Mexico 2016, E) Mediterranean Sea 2013, F) Oregon 2018, G) Oregon 2019, and H) S. California 2010.

